# Efficient strategies to detect genome editing and integrity in CRISPR-Cas9 engineered ESCs

**DOI:** 10.1101/635151

**Authors:** Maja Gehre, Christopher Buccitelli, Nichole Diaz, Jan Korbel, Kyung-Min Noh

**Affiliations:** European Molecular Biology Laboratory (EMBL), Genome Biology Unit, 69117 Heidelberg, Germany; Collaboration for joint PhD degree between EMBL and Heidelberg University, Faculty of Biosciences

## Abstract

CRISPR-mediated genome engineering provides a powerful tool to study the function of genes and proteins. In the past decades, the advances in genome and transcriptome sequencing techniques have shed light on the genetic causes underlying many human diseases, such as neurodevelopmental disabilities or cancer. Sometimes, a single point-mutation in a protein coding gene has been identified as the primary cause of the disease. CRISPR-Cas offers the possibility to introduce or remove such a mutation of interest to understand disease mechanisms and even bears therapeutic potential. We describe the adaptation of an experimental strategy that allows the mutation of protein residues in mouse embryonic stem cells (ESCs) and propose a new screening method, Mismatch-qPCR, to reliably detect editing events in clonal cell lines as an alternative to restriction digest or Sanger sequencing. Finally, we show that RNA-Sequencing (RNA-Seq) data or low-coverage genomic sequencing data can be used to detect large chromosomal deletions and rearrangements that frequently occur at the CRISPR-targeting site.

## Introduction

CRISPR-Cas is an adaptive immune system that protects bacteria and archae against foreign DNA (Jinek et al., 2012; Makarova et al., 2011). In recent years, components of this system have been modified and made applicable for genome engineering in mammalian cells (Charpentier and Doudna, 2013; Ran et al., 2013). The main components are the endonuclease Cas9 that can cleave double-stranded DNA molecules, and a single-guide RNA (sgRNA). The sgRNA acts as a scaffold and directs Cas9 to a genomic site of interest by a short 20 nucleotide complementary guide sequence. The requirement for Cas9 to bind and cleave the targeted genomic sequence is a protospacer adjacent motif (PAM) in the DNA, most commonly a 5’-NGG” motif where N is any nucleotide followed by two guanine nucleotides. Cas9 introduces double-strand breaks into the DNA, which can can be repaired by the Non-homologous end joining (NHEJ) or the homology directed repair (HDR) pathway. NHEJ ligates the DNA strands in an error-prone way that results in insertion or deletion (Indel) mutations at the repair site. Indels can cause frameshift and the formation of premature stop codons, resulting in gene knock-out. The more precise HDR pathway repairs the DNA according to a repair template, which can be a chromosome, exogenously supplied plasmid or single-stranded DNA with homology to the DSB site. This pathways allows precise gene editing by introducing nucleotide changes of interest. Previous studies showed that the frequency of CRISPR-editing via the HDR pathway is low (Ran et al., 2013). Thus, it is required to establish a robust way for high-throughput screening of many clonal cell lines. So far, screening for genome editing is mostly done by restriction digest (Ran et al., 2013) or Sanger sequencing. Restriction digest requires the introduction of nucleotide changes that give rise to a restriction site to help identifying edited clones. This procedure can be tedious and it is not always feasible to change nucleotides in a way to create a restriction site while maintaining the amino acid sequence of the encoded protein. Alternatively, Sanger sequencing is the most reliable technique to help identify editing events in cells, but is expensive if applied to large numbers of clonal cell lines.

In the field of epigenetics, systematic histone mutation studies have already delivered direct clues for the functional significance of histone residues in individual organisms such as yeast and fly (Dai et al., 2008; Hodl and Basler, 2012; Pengelly et al., 2013). In mammalian cells, CRISPR-mediated precise genome editing of histones will help in understanding the role of specific residues and their post-translational modifications for gene regulation in stem cells and during development. Thus, we tested and adapted the CRISPR-Cas9 system developed by Ran et al. (2013) for mutation of histone variant H3.3 in ESCs, specifically to exchange lysine residues with alanine, but our approach can be extended to other genes of interest. For this purpose, we develop a new screening method, Mismatch-qPCR, to reliably detect editing events in clonal cell lines. The new strategy proved to be more time-effective than screening by restriction digest, reduced the costs of screening many cell lines compared to Sanger sequencing and abolished the need to insert a restriction site into the genome. Finally, we address the issue of large chromosomal deletions and rearrangements that have been reported to occur during CRISPR editing (Lee and Kim, 2018; Zhang et al., 2015). For this, we tested the feasibility of leveraging functional genomic data frequently employed in the downstream analysis of CRISPR generated cell lines. Specifically, using sequencing data of either Chromatin Immunoprecipitation inputs (ChIP-Seq) or RNA-Seq data, we detect both large-scale deletions as well as other copy number alterations that occured proximal to the CRISPR-target cut site.

## Results

### Delivery of Cas9 and guide into mouse embryonic stem cells by nucleofection

To target the H3.3 encoding genes *H3f3a* or *H3f3b* for gene editing, we used the plasmid-based CRISPR system developed by Ran et al. (2013). The plasmid encodes a fusion protein of Cas9 and GFP, which allows cell selection by flow cytometry, and a single-guide RNA for targeting of Cas9 to a genomic site. The delivery of this plasmid and a repair template is crucial for successful editing, but primary cells, including ESCs, are difficult to transfect using traditional transduction methods such as liposomal reagents. We tested if electroporation with a nucleofector system (nucleofection) is suitable for the delivery of the CRISPR plasmid into ESCs and analyzed the efficiency by flow cytometry. Overall, around 5% of all sorted cells were GFP-positive and therefore successfully transduced (Fig. 1, gating strategy for this experiment is depicted in Supplementary Fig. S1). This proportion is sufficiently high to obtain the required cell numbers for gene editing. Next, we continued with optimizing the conditions for gene editing of endogenous H3.3 in ESCs using this CRISPR-Cas9 system.

**Figure 1.**
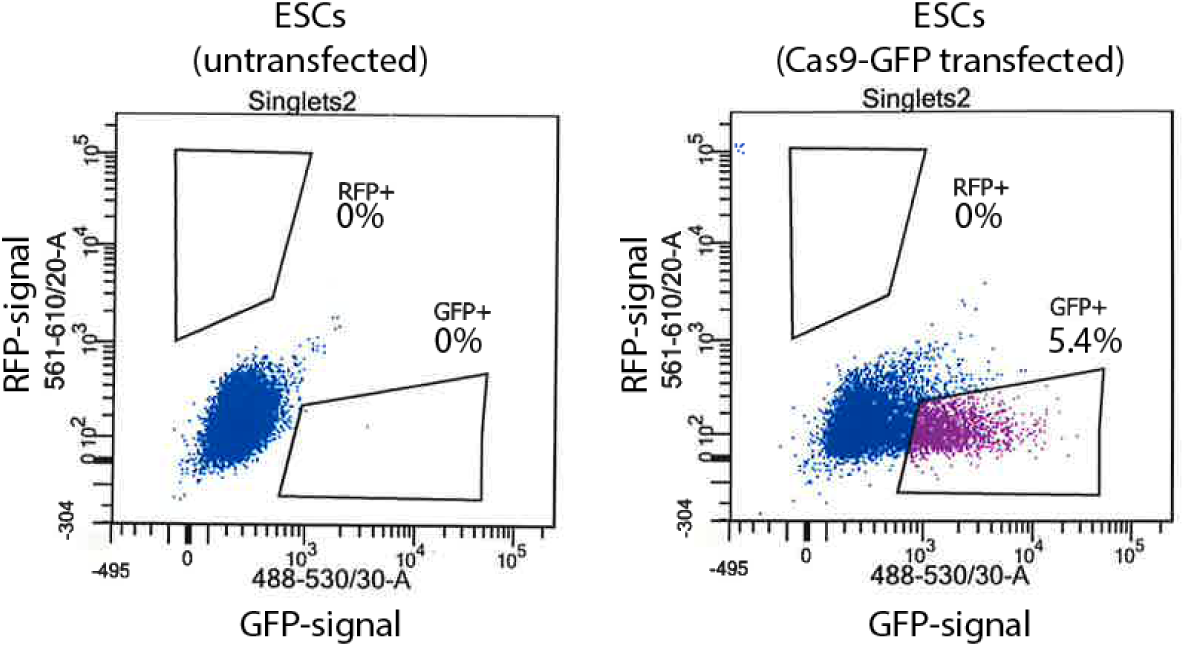
Transduction efficiency in ESCs with a plasmid-based CRISPR system using nucleofection. Nucleofected ESCs were analyzed by flow cytometry for GFP expression, which represents successful delivery of the Cas9-encoding plasmid. Displayed are GFP-signal (x-axis) against RFP-signal (y-axis) and rectangles indicate areas of positive cells expressing the analyzed Cas9-GFP fusion protein.

### Scr7 promoters CRISPR-Cas9 mediated gene editing in ESCs

Gene editing through the HDR pathway occurs at lower frequencies than gene knockout via the NHEJ pathway, but treatment with small molecules has been proposed to promote the frequency of HDR in cells. We tested the efficiency of two small molecules, Scr7 and L755,507, to promote gene editing of the H3.3 genes (*H3f3a* or *H3f3b*). Scr7 has been reported to promote editing via HDR by inhibiting the activity of DNA ligase IV, an important enzyme in the competing NHEJ pathway (Chu et al., 2015; Maruyama et al., 2015). L755,507 is a β3-adrenergic receptor partial agonist reported to enhance gene editing, but the mode of action is unknown (Yu et al., 2015). The treatment with individual small molecules at concentrations between 1-10 μM did not visibly reduce cell viability, but to minimize potential toxicity the cells were treated with the small molecules for only 36 hours of the culture (12 hours before and 24 hours after delivery of the Cas9-plasmid and single-stranded repair template by nucleofection).

In untreated ESCs, we did not obtain edited cell lines carrying the mutation of interest, neither with L755,507 treatment. Using the Scr7 inhibitor, we obtained edited cell lines that had incorporated nucleotide changes according to a supplied repair template. Thus, treatment with Scr7 inhibitor resulted in a higher frequency of editing events than without treatment or by treatment with L755,507 (Table 1). Overall, the editing frequency of either *H3f3a* or *H3f3b* was around 0-2% of transduced cells for homozygous editing and 0-10% for heterozygous editing.

**Table 1.** Examples of CRISPR editing screens to introduce point-mutations into histone H3.3. Non-treated or treated ESCs were transduced with two CRISPR-Cas9 plasmids targeting both the *H3f3a* and *H3f3b* gene. Total number of clonal cell lines that were screened and the detected edited clonal cell lines (heterozygous or homozygous) are indicated.

| Gene | Treatment | Clones | Heterozygous edit | Homozygous edit |
| --- | --- | --- | --- | --- |
| H3f3a | no drug | 56 | 0 | 0 |
| H3f3b | no drug | 56 | 0 | 0 |
| H3f3a | L755,507 | 49 | 0 | 0 |
| H3f3b | L755,507 | 49 | 0 | 0 |
| H3f3a | Scr7 | 43 | 2 (4.7%) | 0 |
| H3f3b | Scr7 | 43 | 0 | 1 (2.3%) |
| H3f3a | Scr7 | 57 | 10 15.8% | 1 (1.8%) |
| H3f3b | Scr7 | 57 | 1 (1.8%) | 1 (1.8%) |

We confirmed the increased frequency of gene editing in the presence of Scr7 in a bulk of Cas9-expressing ESCs by restriction digest. The repair templates carried a restriction site that gets inserted at the repair site in case of successful editing and the efficiency of editing can be estimated by restriction digest in a bulk of cells sorted by flow cytometry. Treatment with Scr7 increased the integration of a restriction site by editing compared to untreated cells (Fig. 2). Despite the treatment of ESCs with an inhibitor to promote HDR, the observed frequencies for precise genome editing were very low. The editing frequency we observed in the initial experiments for homozygous editing of *H3f3b* were approximately 1.5% and it would therefore require screening of many clonal cell lines to obtain a successfully edited clone per mutation.

**Figure 2.**
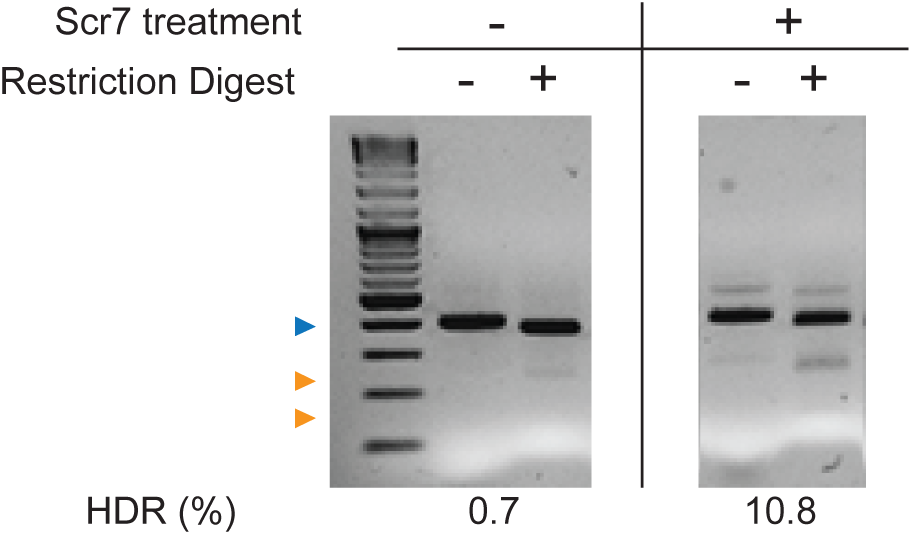
Scr7 promotes CRISPR gene editing in ESCs. Cells were transduced with CRISPR-Cas9 plasmids targeting the *H3f3b* gene and repair templates carrying a restriction site for genomic insertion. A bulk of cells expressing Cas9-GFP was selected by flow cytometry. The targeted *H3f3b* locus was amplified by PCR and subjected to restriction digest. Digestion pattern was analyzed by agarose gel electrophoresis. Successful integration of the restriction site results in cleavage of the PCR product (blue arrowhead) and the occurrence of smaller digestion products (orange arrowheads). HDR frequency was calculated as the ratio of band intensities.

### Mismatch-qPCR as a high-throughput screening method to detect gene editing

The systematic exchange of multiple protein residues of in mammalian cells can only be achieved if a reliable method allows high-throughout screening of many clonal cell lines at reduced costs. Whereas Sanger sequencing is a fast and precise method for screening, it is expensive if applied to many cell lines. Instead, screening by restriction digest is inexpensive, but tedious and requires the insertion of a restriction site into the targeted genomic locus.

We tested if CRISPR-editing events can be detected in a quantitative PCR (qPCR) by designing mutation-specific primers that recognize the inserted nucleotide changes of interest (Fig. 3a,b), referred to as Mismatch-qPCR.

**Figure 3.**
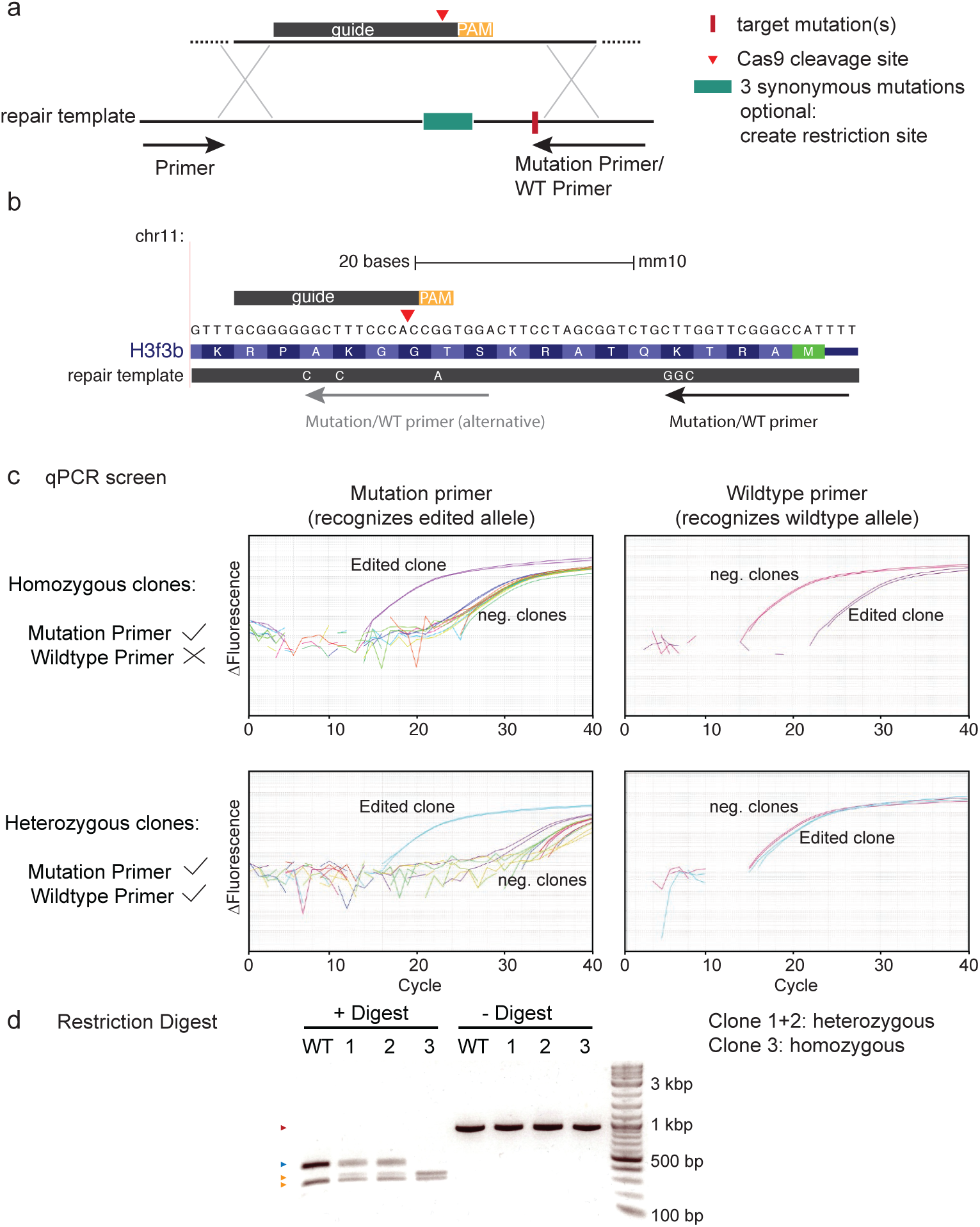
Mismatch qPCR screen detects CRISPR-mediated point-mutations in H3.3B. (a) Guide sequences were designed to direct Cas9 close to the mutation site of interest. The repair template contains nucleotide changes to introduce a target mutation, and 3 additional synonymous mutations into the guide binding site or PAM to prevent re-cleavage after repair. Optionally, synonymous mutations can give rise to a new restriction site used to validate clones. The mutation-specific primer recognizes nucleotide changes that arise after CRISPR-editing at the most 3’end. The wild-type (WT) primer recognizes the same, but unmodified genomic site. (b) Examples of two mutation-specific primers for Mismatch qPCR that detect editing of lysine 4 to alanine in H3.3B by recognizing either the K4A mutation or the synonymous mutations inside the guide. (c) Mismatch qPCR screen of CRISPR cell lines using mutation-specific and wild-type primers. Successful amplification result in an increase of the fluorescent signal (y-axis) at lower cycle numbers (x-axis). DNA of homozygously edited clones is amplified only with mutation-specific primers, whereas heterozygous clones are also amplified using the wild-type primer. (d) Confirmation of editing events by restriction digest using a newly introduced restriction site after CRISPR targeting. DNA of wildtype cells (WT) and positive clones predicted by Mismatch qPCR screening were used for PCR amplification followed by restriction digest with BanI. Digestion pattern was analyzed by agarose gel electrophoresis. Digestion of the PCR product (red arrowhead) of wild type DNA results in a larger product (blue arrowhead) than from edited DNA (orange arrowheads) with an additional integrated restriction site. Restriction digestion confirms the detected editing events by qPCR.

Using this method, we were able to separate edited clones from wild type clones by shifts to lower cycle threshold numbers (rounds of amplification) (Fig. 3c). In combination with a primer that recognizes the unchanged wild type allele, it was possible to distinguish heterozygous from homozygous clones. Heterozygous clones with one mutant and one wild type allele amplify in a qPCR reaction with both primer sets, while homozygous clones only amplify using the mutation-specific primer. Using restriction digest, we confirmed homozygosity and heterozygosity of the clonal lines, which can be identified by the complete or incomplete digestion of a PCR product (Fig. 3d) and the results were in agreement with the results from the qPCR screen. After identification of potential candidate clones by Mismatch-qPCR, the exact genotype of the edited clones has to be determined by Sanger sequencing to confirm that the mutation was introduced correctly in both alleles. Hereby, we confirmed the successful exchange of lysine 4 or 36 in H3.3B (Fig. S2). For some candidate clones that were detected by screening, we observed incomplete repair resulting in additional small deletions around the guide binding site. Only clonal lines that have incorporated nucleotide changes correctly from the repair template can be used for downstream analysis.

### CRISPR off-target analysis for gene copy number alterations

Double-strand cleavage by Cas9 can cause unintended off-target effects that affect genome integrity (Lee and Kim, 2018; Zhang et al., 2015). As a next step, we wanted to confirm that during clonal selection and CRISPR targeting the integrity of the genome was not affected in selected clones, e.g. by chromosomal rearrangements.

Gene expression data has been previously demonstrated to be predictive of somatic gene copy number alterations in the absence of accompanying genomic data in cancer cells (Ben-David; et al., 2016; Fehrmann et al., 2015). Since gene expression data is regularly generated in contemporary genomic studies from CRISPR-edited or knock-out cell lines, we tested if it can be exploited to confirm their genomic integrity after targeting. The advantage would be to exclude clonal cell lines with chromosomal deletions or duplications, which could otherwise complicate downstream analysis. Using mRNA-Seq, we determined gene expression changes in CRISPR cell lines relative to their wild type ESC line of the same genetic background. The gene expression changes were compared to the genomic coordinates of the respective gene. A sequence of down- or up-regulated genes that are located in proximity to each other indicates large-scale chromosomal abnormalities (Fig. 4a). Using this strategy, we found that incomplete repair of a chromosome can result in large copy number alterations (frequently chromosome arm losses), typically beginning at the CRISPR target site and spanning the rest of the chromosome arm (Fig. 4b,c,d). Such events result in hemizygous loss of hundreds of genes. We observed that chromosome arm-losses can occur independently of the exact CRISPR guide sequence and on different chromosomes, given that they were detected during the targeting of *H3f3a* on chromosome 1 or *H3f3b* on chromosome 11 (Fig. 4b,c). Both H3.3-encoding genes are located at the periphery of chromosome 1 and 11, respectively, and it is possible that CRISPR-targeting of genes at the ends of chromosomes are more likely to result in hemizygous chromosome-arm loss since larger changes in gene copy number that would result from a mid-chromosome cut may be less well tolerated. It should be noted that such deletions are not detectable by traditional Sanger Sequencing, because only the intact allele is amplified in a PCR reaction. Additionally, we also observed rearrangements of chromosomes that were not targeted by CRISPR, e.g. of chromosome 6 (Fig. 4d). These rearrangements can potentially be CRISPR off-target effect, but may also have occurred spontaneously during clonal selection.

**Figure 4.**
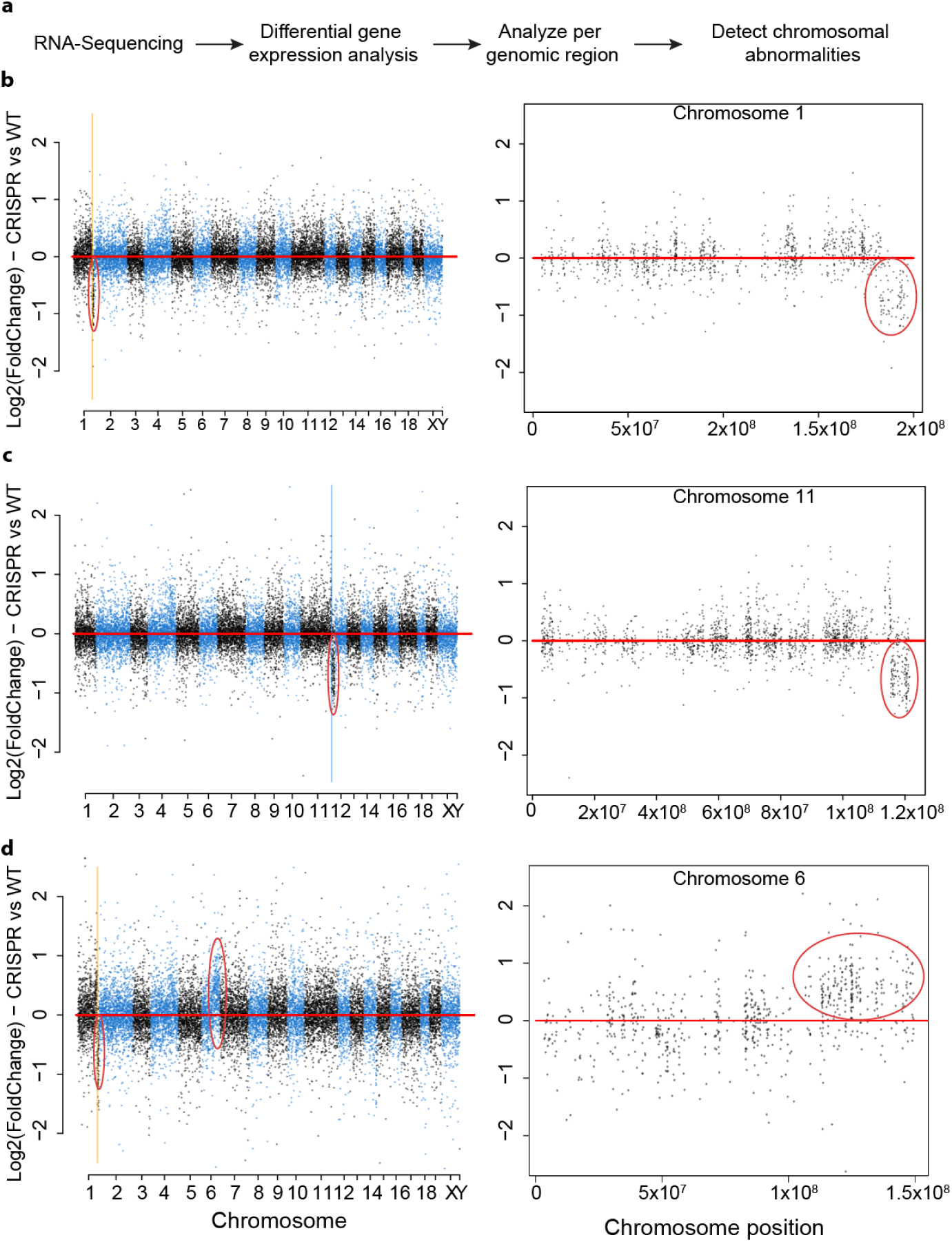
Prediction of chromosomal rearrangements from RNA-Sequencing data. (a) Strategy for analyzing genome integrity from RNA-Seq data. (b,c,d) CRISPR off-target analysis from differentially expressed genes. log2(FoldChanges) of gene expression in three different CRISPR clones compared to unmodified wild type ESCs were determined by DESeq2 and plotted over chromosome position for all (left) or a specific chromosome (right). Lines indicate CIRSPR cleavage site inside *H3f3a* gene (yellow) and *H3f3b* gene (blue). Loss or duplication of a chromosome part can be detected by coordinated up- or down-regulation of proximal genes. (b) CRISPR clone showing a cluster of systematically down-regulated genes on chromosome 1 close to the CRISPR targeting site in *H3f3a* gene. (c) CRISPR clone showing a cluster of down-regulated genes on chromosome 11 close to the CRISPR targeting site in *H3f3b* gene. (d) CRISPR clone showing a cluster of down-regulated genes on chromosome 1 close to the CRISPR targeting site in *H3f3a* gene and additionally a large cluster of up-regulated genes on chromosome 6 (likely a consequence of clonal selection and unrelated to CRISPR targeting).

To increase the confidence in genome integrity predictions from RNA-Seq data, we tested whether the predicted chromosomal deletions/duplications can be confirmed on the genomic level by using low-coverage genomic sequencing data, such as ChIP-Seq Inputs. From RNA-Seq data of a chosen cell line, a cluster of up-regulated genes and a cluster of down-regulated genes between the CRISPR target site and the chromosome end were detected (Fig. 5a), perhaps indicative of a simple breakage fusion bridge cycle initiated by a DNA break (Bignell et al., 2007). The same chromosomal abnormality was also detectable using ChIP-Seq Input data (Fig. 5b), which confirmed that the gene expression changes were the result of a duplication-deletion rearrangement on the genomic level. Compared to the genomic analysis using ChIP-Seq Input data, RNA-Seq data yields a lower resolution because the predictions are dependent on the gene-density per chromosome, which is rather sparse considering that only 62% of the genome is transcribed, and an even smaller fraction of this corresponds to coding exons (5.5%) (Consortium, 2012). Thus, low-coverage genomic sequencing data (e.g. ChIP-Seq) allows a more detailed analysis of chromosomal abnormalities with higher precision and confidence, but with RNA-Seq data it is possible to make similar predictions especially in the case of large chromosomal abnormalities.

**Figure 5.**
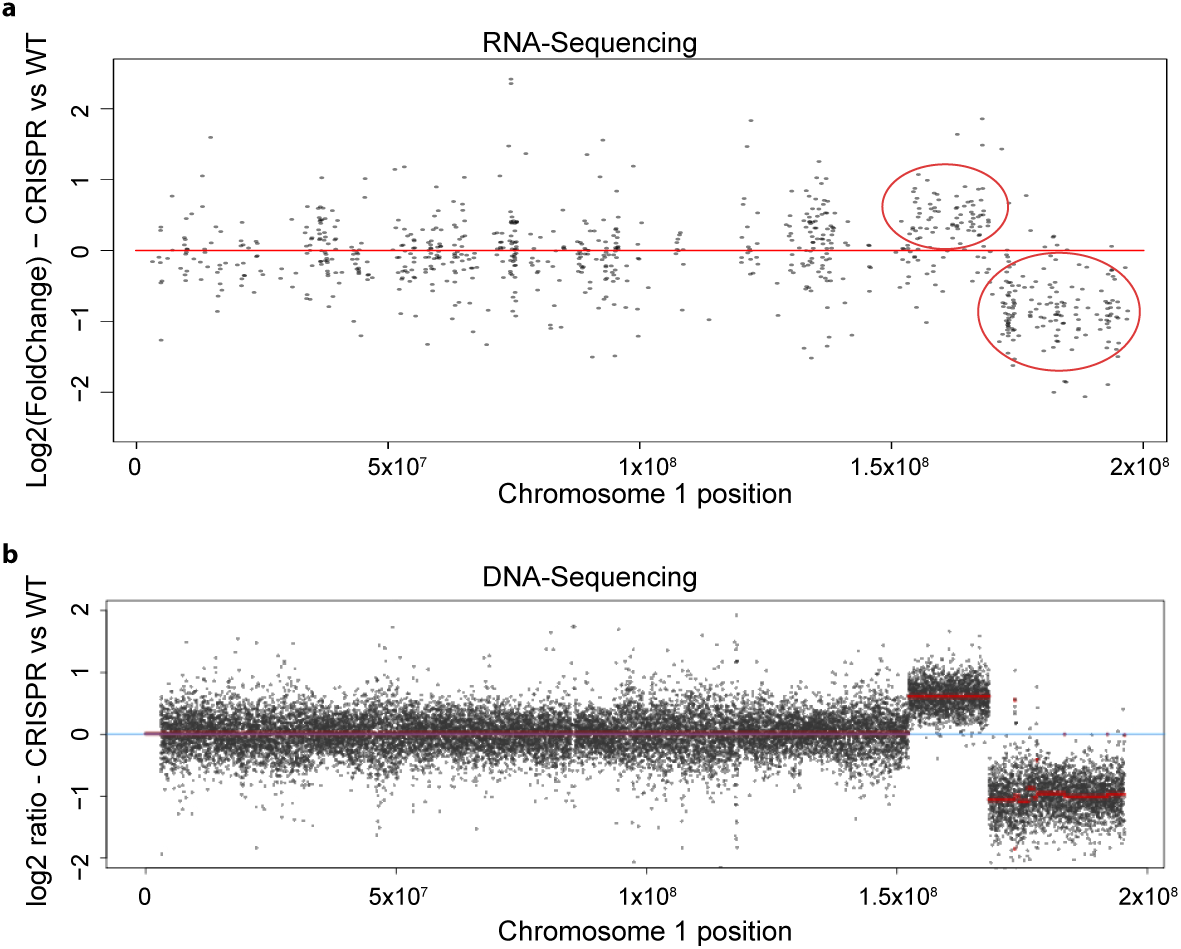
Comparison between RNA-Seq and ChIP-Seq Input data in predicting chromosomal rearrangements. (a) Differential gene expression analysis with regard to the genomic coordinates from RNA-Seq data. CRISPR clone shows two clusters of systematically up- and down-regulated genes on chromosome 1 close to the CRISPR targeting site, suggesting a partial chromosomal duplication and deletion. (b) Analysis of ChIP-Seq Input reads with respect to genomic coordinates confirms the duplication-deletion rearrangement at the CRISPR targeting site on chromosome 1 on the genomic level.

The occurrence of CRISPR-dependent and -independent effects on genome integrity suggests that an extensive on- and off-targeted analysis for generated clonal cell lines is recommended and should be integrated into the standard workflow for validation of CRISPR cell lines. The summarized strategy for gene editing in ESCs was developed for editing of H3.3 encoding genes in mouse ESCs, but has successfully been applied to other genes of interest (Fig. 6)

**Figure 6.**
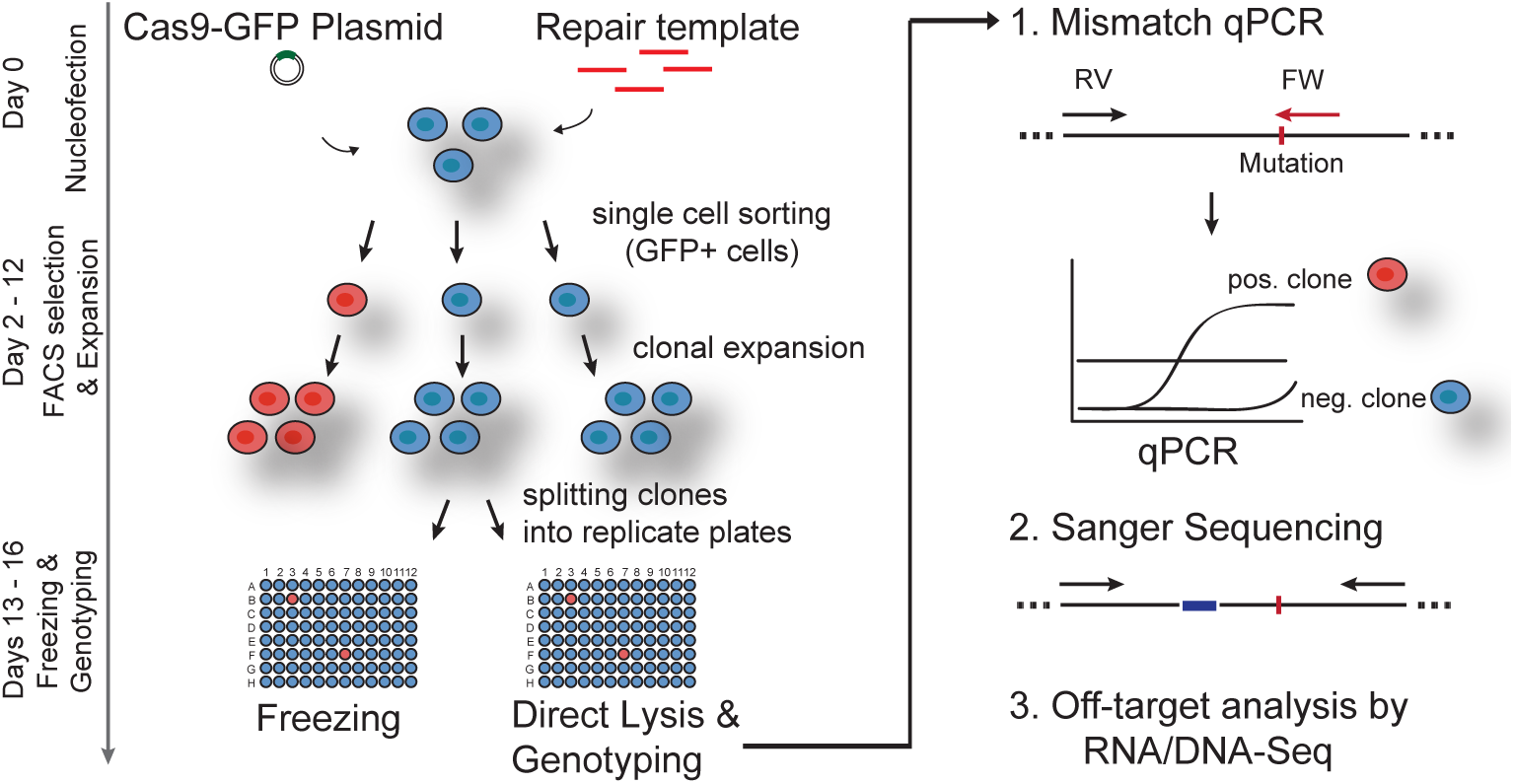
Overview of CRISPR-editing workflow in mouse ESCs. General scheme used to introduce point-mutations into H3.3B. Cas-9 plasmid with guide and single-stranded repair templates are delivered into ESCs by nucleofection. Transduced GFP-positive cells are selected by flow cytometry and single cells are sorted into 96-well plates. Clonal cell lines are expanded and split into two identical plates for freezing and screening. For Mismatch-qPCR, cells are lysed directly in a 96-well plate and successful editing events are detected in a qPCR reaction. Editing events can be confirmed by Sanger sequencing and optionally by restriction digest. Large-scale on- and off-target analysis is performed in selected clonal lines from RNA- or DNA-Seq data.

## Discussion

Genome engineering by CRISPR-Cas9 provides a powerful tool to exchange endogenous protein residues and to probe their function in mammalian development. Targeting of the H3.3 genes (*H3f3a* and *H3f3b*) showed that gene editing occured at low frequencies in mouse ESCs and it required extensive screening of many clonal cell lines to obtain successfully edited clones. To screen clonal cell lines with high through-put, we developed a qPCR-based method that could reliably identify CRISPR-edited clones. By direct comparison with the restriction digest method (Ran et al., 2013), Mismatch-qPCR proved to be a faster screening method and did not require the insertion of a restriction site into the genome. The read-out can be observed during the qPCR reaction without requiring subsequent analysis steps. Sanger sequencing was required to exclude false positive clones and to confirm the precise genotype of the clonal cell lines.

Nevertheless, sequencing of few candidate clones after screening by Mismatch-qPCR was more economical than to sequence all generated clonal cell lines. The limitation of this approach is certainly the requirement of suitable primer pairs for screening. Dependent on the DNA sequence and GC-content of the targeted locus, it is not always possible to design primers that fall into the recommended property range (e.g. melting temperature), which is predicted to result in less efficient PCR amplification. However, the read-out of Mismatch-qPCR is qualitative and not quantitative and should not necessarily be compromised by less efficient primers.

Following the generation on CRISPR-edited clones, we wanted to confirm that their genomic integrity had not been compromised by the targeting. Chromosomal rearrangements can occur simply during clonal selection of ESCs and large chromosomal deletions and complex genomic rearrangements can occur at the CRISPR-targeted site in mouse ESCs as recent studies have suggested (Kosicki et al., 2018). Using RNA-Seq data, we frequently observed deletions at the site of CRISPR-targeting, and less commonly rearrangements on other chromosomes. At the CRISPR-target site, double-strand DNA cleavage resulted in the one-allelic loss of a chromosome arm and thus down-regulation of hundreds of genes. These results caution against neglecting the risk of on- and off-target effects introduced by CRISPR-Cas9. Testing the genomic integrity of generated clonal lines from commonly available genomic datasets such as RNA-Seq and low-coverage DNA-Seq data (e.g. 1x coverage) can help in excluding affected cell lines and indeed, if not available, other -omics data that scales with copy number (such as proteomics) may also be explored for this purpose.

Large-scale deletions and rearrangements severely affected genome integrity, and suggest that an extensive on- and off-target analysis for generated clonal cell lines is indeed necessary and should be integrated into the standard workflow for CRISPR editing. Depending on the availability, both RNA- or DNA-Sequencing data are suitable for this analysis and these datasets are often available for already published studies, e.g. input DNA sequencing data from ChIP experiments. A sequencing based approach for identifying off-target clones will be easily implemented and might be comparable with the traditional karyotype analysis.

With the growing list of mutations associated with human diseases, CRISPR-Cas9 mediated editing is becoming increasingly important to study disease mechanisms. Economical screening methods with high-throughput such as Mismatch qPCR in combination with a large-scale off-target analysis can facilitate the generation of multiple biological replicates for a mutation, which is essential for data interpretation and reproducibility.

## Materials and Methods

### Cell culture

Murine ESCs (129XC57BL/6J) were cultured in ESC media containing Knockout-DMEM (Thermo Fisher) with 15% EmbryoMax FBS (Millipore) and 20 ng/ml leukemia inhibitory factor (LIF, produced by Protein Expression Facility at EMBL Heidelberg), 1% non-essential amino acids, 1% Glutamax, 1% Pen/Strep, 1% of 55mM beta-Mercaptoethanol solution. Cells were maintained at 37°C with 5% CO2. ESCs were routinely tested for mycoplasma.

### Guide design and CRISPR plasmid cloning

Guides were designed with homology to a sequence close to the mutation site of interest using MIT’s Optimized CRISPR design tool. As a general guideline, the guide binding site should ideally be less than 30 nucleotides away from the mutation site of interest, and can also overlap the mutation site. If the mutation site is close to an intron, it is recommendend to use an intronic guide sequence in case additional indels occur at the CRISPR cutting site, but this is optional. Guide sequences with an aggregate score of greater than 50% were selected and cloned into pSpCas9(BB)-2A-GFP (PX458, Addgene) or pSpCas9(BB)-2A-RFP (modified from PX458) according to instructions by Ran et al. (2013). For this purpose, phosphorylated DNA oligos (5’-Phos) were ordered from Eurofins according to this scheme:

CACC + G + guide sequence forward

AAAC + guide sequence reverse + C

pSpCas9(BB)-2A-GFP was digested with BbsI, followed by dephosphorylation using Antarctica Phosphatase (NEB) and separated from undigested plasmid by 1% agarose gel electrophoresis. Digested plasmid was extracted from the gel (Gel extraction Kit, Qiagen). Complementary guide oligos were annealed and cloned into pSpCas9(BB)-2A-GFP/RFP plasmid, from here on referred to as Cas9-GFP-guide or Cas9-RFP-guide plasmid.

We used guides with the following sequences:

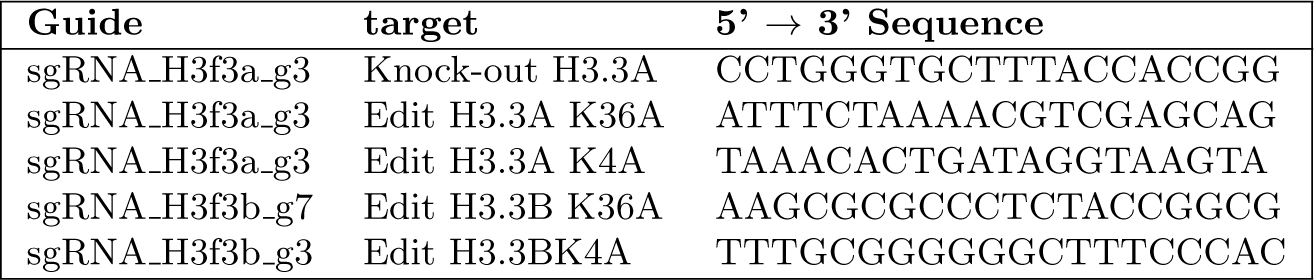

### Design of repair template

The single-stranded repair template was designed to encompass the DNA 90 bp upstream and downstream of the Cas9 cutting site as suggested by Ran et al. (2013). We used IDT’s DNA ultrameres (up to 200 bp) as DNA template. The repair template contains the mutation of interest (e.g. a lysine to alanine exchange in H3.3 at lysine 4 or lysine 36) and 3 synonymous mutations inside the guide binding site to prevent repeated cleavage by Cas9. Editing efficiency can be improved if one of these additional mutations changes the PAM sequence into a non-PAM sequence. Synonymous mutations do not change the resulting protein sequence, and should be chosen in a way that codon usage frequency is considered and codons with very low frequencies should not be used as they can alter expression levels of the encoded protein. We used single-stranded DNA templates with the following sequences:

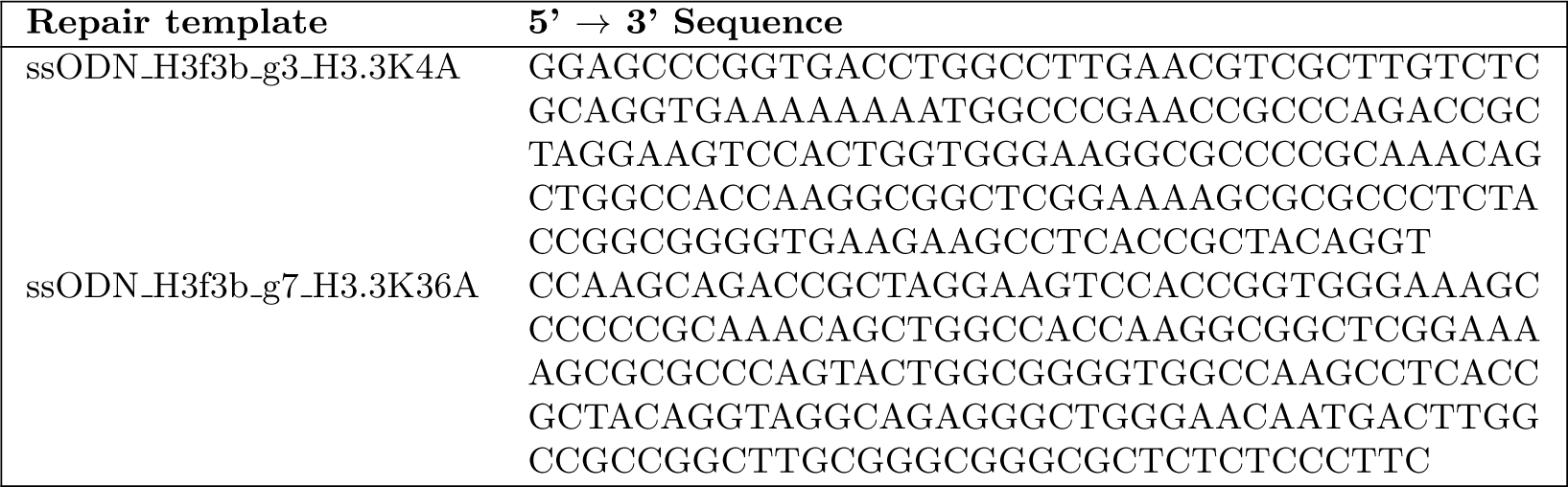

### Nucleofection

For gene editing 2×10^6^ ESCs were transfected with 2 μg Cas9-GFP-guide plasmid and 5 μl of 100 μM ssODN repair templates (180 bp, IDT ultrameres) using electroporation (Nucleofector, Lonza). Cells were resuspended in 100 μl of P3 solution (Lonza) and 2 μg plasmid DNA and 5 μl donor template were added. Cells were transferred into a cuvette and electroporated with pulse code CG 104 for mouse ESCs (“ES, mouse“). After electroporation cells were plated into a T-25s flask with pre-plated MEFs containing fresh ESC media. For drug treatment of ESCs, media was supplemented with 5 μM L755,505 (Xcessbio, M60237-2s) or 10 μM Scr7 inhibitor (Xcessbio, M60082-2s) for 12 hours prior to nucleofection and for additional 24 hours after nucleofection to promote gene editing (Chu et al., 2015; Maruyama et al., 2015; Yu et al., 2015).

### Single-cell sorting by flow cytometry

Single-cell sorting by flow cytometry was performed 48 hours post-nucleofection for GFP positive cells. Single cells were sorted into the wells of multiple 96-well plates containing pre-plated mouse embryonic fibroblasts (MEFs) as feeders and 150 μl of ESC media per well. In one well, a bulk of 1.000-10.000 cells was sorted serving as a positive control during screening. Cells were sorted on a FacsAria Fusion sorter (BD Biosciences). Gating was performed with BD’s FACSDiva 8.0.1 software and single cells were chosen for analysis after doublet discrimination. The gating strategy for GFP-positive transfected cells is depicted in Supp. Fig. S1.

### Cell expansion, freezing and lysis of clonal lines

Growing ESC colonies were dispersed 6 days after sorting by trypsinization. After disperal, ESC media was changed every day in wells with growing ESC colonies. Dispersed clones were split 9 days after sorting into 2 replicate 96-well plates. One plate was used for freezing in DMSO-containing medium and the second was used for lysis and screening of clones. To detect gene editing events, clonal cell lines were lysed directly in one of the replicate 96-well plates. For cell lysis medium was removed from wells and 70 μl of lysis buffer were added to each well. Lysis buffer was prepared by diluting one part direct-lysis reagent (301-C, Viagen Biotech) in two parts destilled water (e.g. 100 μl buffer and 200 μl H2O) and supplemented with 0.5 mg/ml proteinase K (03115887001, Roche). Cells were lysed at 55°C for 2 hours while shaking at 350 rpm, and afterwards proteinase K was heat deactivated for 45 min at 85°C. Cell lysates can be stored at 4°C for up to one week, for long-time storage (1-2 months) lysates were frozen at −20°C. To freeze cell clones in replicate plate, prepare a new 96-well round-bottom plate (168136, Thermo Fisher) with 50 μl of freezing media (10% Knock-Out DMEM, 60% FCS, 30% DMSO). Remove media from wells in second replicate plate and using a multichannel pipette wash cells with PBS and add 35 μl of trypsin per well, trypsinize cells for 5 minutes at 37°C. Quench trypsin with 65 μl of ESC media. Resuspend cells by pipetting using a multichannel pipette and transfer cells drop by drop to round-bottom plate containing freezing media, gently pipette up and down to mix. After transfering all wells, wrap plate with parafilm and paper towels and place plates in Styrofoam box for freezing at −80°C.

### Mismatch-qPCR

Screening primers to detect genome editing events by qPCR should be suitable for standard qPCR reaction and the total amplicon size should be under 150 bp to guarantee successful amplification during elongation step. One of the primers encompasses the editing site, ideally directly ending with a point mutation on the 3’ end. Thus, amplification with this primer should not work on the wild type sequence. Instead, the wild-type primer is designed to recognize the unmodified genomic sequence. Reverse primer recognizes the wild-type sequence away from the CRISPR editing site. For editing screen, 0.5 μl of the crude lysate are sufficient for a 20 μl qPCR reaction (96-well plate) or 0.25 μl for a 10 μl qPCR reaction (384-well plate), higher amounts of crude lysate can interfere with PCR reaction. DNA from a bulk of sorted cells (1.000-10.000 cells) should be included as a positive control and wild type DNA from unmodified cells as a negative control. qPCR is run with a standard cycling program to detect cycle threshold (Ct) values. All clones with significantly lower Ct numbers than the negative control (wild type), and similar or lower Ct values than the positive control were used for downstream validation. For Sanger sequencing, a region of 1-1.5 kb around the mutation site was amplified by PCR and sequenced from both ends to confirm editing. The crude lysates can also be used for genotyping, but may result in lower quality of Sanger sequencing results. In this case it is recommended to thaw all positive clones, expand them and extract genomic DNA (Puregene Core Kit B, Qiagen) to improve the quality of the sequencing reaction. Editing events were confirmed and checked for homozygosity by analyzing the chromatogram (SnapGene Viewer) of the Sanger Sequencing reaction.

### Restriction Digest

A DNA fragment of 1kb around the CRISPR cutting site was amplified using PCR. PCR product was digested with a suitable restriction enzyme (for which restriction site was inserted into the genome) for 30 minutes at 37°C. Digestion products were analyzed by agarose gel electrophoresis.

### CRISPR off-target analysis by RNA-Seq analysis

CRISPR off-target effects in the form of chromosomal duplications/deletions were ruled out by RNA-sequencing. RNAs were extracted from approx. 1×10^6^ cells using RNeasy Kit (Qiagen), followed by DNase digestion using TURBO DNase (Thermo Fisher). mRNAs were isolated from 1 μg of total RNA using a PolyA selection kit (NEB) and sequencing libraries were prepared following instructions from NEBs Ultra Library Preparation Kit for Illumina. All samples were barcoded, pooled and sequenced on a HiSeq2000 Sequencer (Illumina) using a 50 bp single-end run. Sequencing reads were mapped to mouse reference genome (mm10 assembly) using Tophat2 aligner with default settings for single-end reads. Reads per gene were counted using HTSeqCount union or intersection nonempty mode. We used Ensembl gene annotation Mus musculus.GRCm38.83. Differential RNA-Seq analysis between each clone and wild type cells was performed using DESeq2 package (Bioconductor, (Love et al., 2014)) to obtain log2(FoldChanges) per gene. Genomic coordinates per gene were obtained using Biomart (Bioconductor) and log2(FoldChanges) were plotted over chromosome position to obtain distribution profiles of gene expression changes. Cell lines that displayed deletions or duplications of chromosome regions, as seen by concomitant up- or down-regulation of close-by genes, were discarded and not used for analysis.

### CRISPR off-target analysis by RNA-Seq analysis

Purified DNA was fragmented into 500 bp fragments by sonication using Bioruptor Pico (Diagenode). Sequencing libraries were prepared using DNA Ultra II library preparation kit (NEB) according to the manufacturer’s instructions and sequenced either on Illumina’s HiSeq2000 Sequencer (50 bp single-end mode) or NextSeq 500 Sequencer (75 bp single-end mode). Sequencing reads were aligned to mouse reference genome (mm10 assembly) using Bowtie2 (Langmead and Salzberg, 2012). Only non-duplicated, uniquely mapped reads were retained for further analysis.

## Supplementary Material

**Figure S1.**
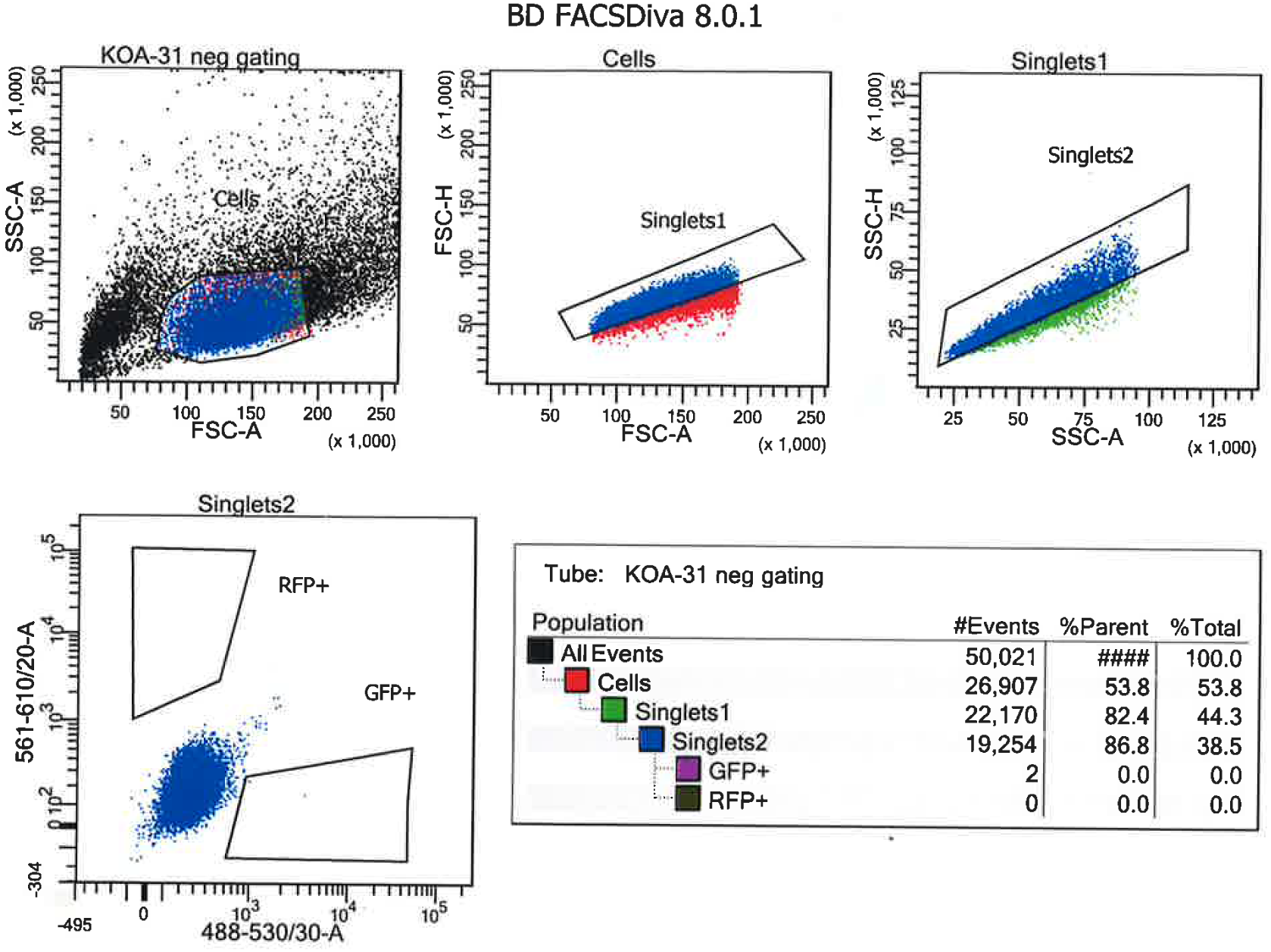
Gating strategy for Cas9-GFP-positive cells flow cytometry analysis. Representative gating strategy for an untransfected wild type cells is displayed. Single cells were chosen for analysis after doublet discrimination by detection of disproportions between cell size (FSC-A) vs. cell signal (FSC-H). The same cell displays higher correlation on the two axis (FSC-A/SSC-A and FSC-H/SSC-H) and all singlet events will fall more on a diagonal than doublets. Transduced GFP-positive cells can be detected outside of the negative population of cells measured with a 488-530 nm laser.

**Figure S2.**
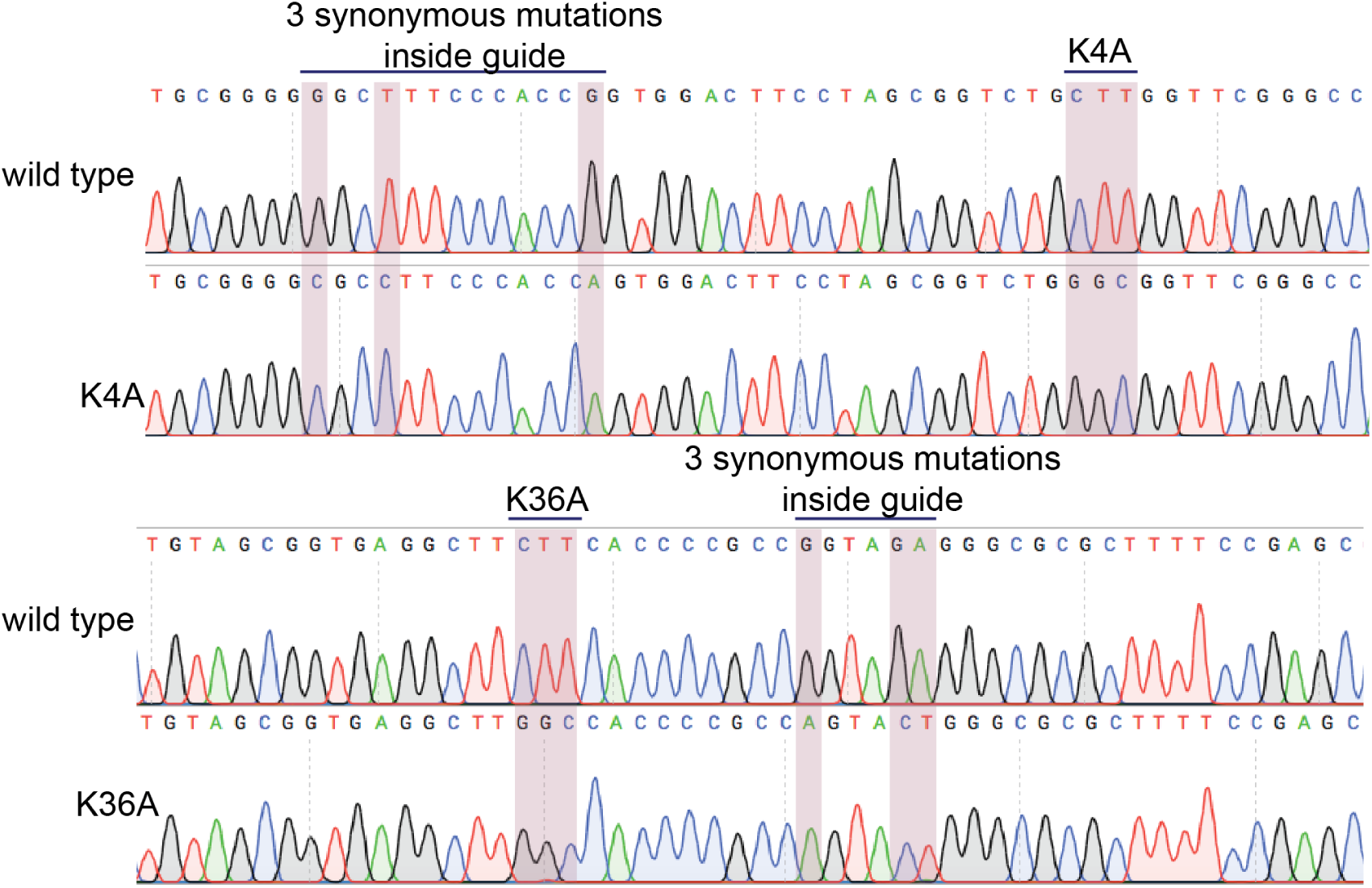
Sanger sequencing confirming the successful integration of nucleotide changes into H3.3B at lysine 4 and 36. (a) Sanger-sequencing results of the H3.3B locus for H3.3K4A and H3.3K36A mutant cells. Analysis of chromatograms from Sanger sequencing confirms the homozygous exchange of targeted nucleotides in H3.3K4A/K36A mutant cell lines. Lysine-to-alanine mutation at either K4 or K36, respectively, and the introduction of 3 additional synonymous mutations inside guide recognition site or PAM are indicated.

## Supporting Information

## Acknowledgments

We thank the staff at the European Molecular Biology Laboratory (EMBL) Genomics Core facility, the Flow Cytometry Core Facility, the Genome Biology Computational Support and the members of the Noh lab for helpful discussions.

